# Replaying Evolution to Test the Cause of Extinction of One Ecotype in an Experimentally Evolved Population

**DOI:** 10.1101/022798

**Authors:** Caroline B. Turner, Zachary D. Blount, Richard E. Lenski

## Abstract

In a long-term evolution experiment with *Escherichia coli*, bacteria in one of twelve populations evolved the ability to consume citrate, a previously unexploited resource in a glucose-limited medium. This innovation led to the frequency-dependent coexistence of citrate-consuming (Cit^+^) and non-consuming (Cit^−^) ecotypes, with Cit^−^ bacteria persisting on the exogenously supplied glucose as well as other carbon molecules released by the Cit^+^ bacteria. After more than 10,000 generations of coexistence, however, the Cit^−^ lineage went extinct; cells with the Cit^−^ phenotype dropped to levels below detection, and the Cit^−^ clade could not be detected by molecular assays based on its unique genotype. We hypothesized that this extinction event was a deterministic outcome of evolutionary change within the population, specifically the appearance of a more-fit Cit^+^ ecotype that competitively excluded the Cit^−^ ecotype. We tested this hypothesis by re-evolving the population from one frozen sample taken just prior to the extinction and from another sample taken several thousand generations earlier, in each case for 500 generations and with 20-fold replication. To our surprise, the Cit^−^ type did not go extinct in any of these replays, and Cit^−^ cells also persisted in a single replicate that was propagated for 3,000 generations. Even more unexpectedly, we showed that the Cit^−^ ecotype could reinvade the Cit^+^ population after its extinction. Taken together, these results indicate that the extinction of the Cit^−^ ecotype was not a deterministic outcome driven by competitive exclusion by the Cit^+^ ecotype. The extinction also cannot be explained by demographic stochasticity, as the population size of the Cit^−^ ecotype should have been many thousands of cells even during the daily transfer events. Instead, we infer that the extinction must have been caused by a rare chance event in which some aspect of the experimental conditions was inadvertently perturbed.

## Introduction

In studying patterns of extinction over the course of evolutionary history, a key question is the extent to which extinctions are random or deterministic events [1, 2]. Very small populations can go extinct as a consequence of demographic stochasticity [3-5], that is, random fluctuations in population size that reflect the discrete nature of birth and death processes in finite populations. In the case of sexually reproducing organisms, inbreeding depression can also contribute to the extinction of small populations [6, 7]. Other causes of extinction, such as asteroid impacts and anthropogenic changes, may also be random (at least insofar as they are exogenous to the affected organisms), yet they can eliminate large populations and even extinguish entire clades [1, 8]. In other cases, though, extinctions might be more or less deterministic in the following sense: if the extinctions reflect processes that are endogenous to the organisms and the communities in which they are embedded, then the extinctions would be predictable and repeatable if the same system could be observed multiple times. For example, one lineage may be fated to extinction because it cannot evolve and adapt as well as its competitors or other biological enemies [9, 10]. These different causes of extinction may also have different consequences for how an ecosystem recovers from extinction [11, 12].

Unfortunately, it is difficult to identify definitively the causes of most extinctions in geological history. Even for contemporary extinctions, one cannot generally manipulate and test possible causes of extinction by taking the extinct organisms and reintroducing them into the same ecosystems. However, as we will show, when an extinction occurs during a controlled laboratory experiment with microbes, it is possible, at least in principle, to perform replay experiments to test certain causes of extinction directly and determine whether the extinction was a random or deterministic event.

Here we examine possible causes of an extinction event in a long-term evolution experiment with *E. coli* (LTEE). Because *E. coli* can be revived from frozen stocks [13], the LTEE has a living fossil record with which a population’s evolution can be replayed, starting from one or more frozen samples, under conditions that are as close as possible to those of the original evolutionary event [10, 14]. In the event of a lineage’s extinction, these samples allow the population to be revived from a point prior to the extinction and re-evolved in many parallel replicates to determine the frequency with which that lineage’s extinction happens again. Microbes have the additional advantages of short generation times and large population sizes, allowing observation of substantial evolutionary change over tractable time spans [13, 15].

The LTEE consists of 12 independently evolving populations, each of which was founded from one of two *E. coli* strains that differ by a selectively neutral marker [13]. Growth in the LTEE is carbon-limited, with glucose as the primary carbon source. The 12 populations had been evolving for more than 50,000 generations at the time of the experiments reported here. In one of the 12 populations (called Ara-3), the novel ability to grow aerobically on citrate evolved after more than 30,000 generations [14]. The inability to grow aerobically on citrate is one of the identifying traits of *E. coli* [16], so the abundant citrate present in the LTEE culture medium was previously unavailable to the bacteria. The population size increased several-fold after citrate consumption evolved [14], and the evolution of that new ability substantially altered the ecological conditions of the population [17].

The citrate-consuming lineage (Cit^+^) did not fix in the population but instead coexisted with a clade that could not grow on citrate (Cit^−^). These two clades coexisted in a negative frequency-dependent fashion for more than 10,000 generations. The initial phase of coexistence involved a tradeoff, whereby the Cit^+^ cells grew more slowly on glucose than their Cit^−^ counterparts during part of the daily growth cycle (Blount et al. 2008). In particular, the Cit^+^ cells exhibited a longer lag phase prior to commencing growth after the daily transfers into fresh medium. Following the emergence of the Cit^+^ lineage, each clade continued to evolve, though with a different suite of mutations accumulating in each [18]. Notably, the Cit^+^ clade evolved a higher mutation rate and accumulated many more mutations than did the Cit^−^ clade [18]. The Cit^−^ cells adapted to the presence of the Cit^+^ lineage, at least in part, by evolving the ability to consume additional energy-containing carbon molecules released by the Cit^+^ bacteria [17]. The physiological mechanism for the evolution of citrate consumption in the Cit^+^ lineage was a mutation causing aerobic expression of the CitT transporter protein [18]. CitT is an antiporter that allows the Cit^+^ bacteria to import citrate in exchange for various C_4_-dicarboxylic acids including succinate, malate, and fumarate [19]. The Cit^−^ ecotype evolved improved growth on these C_4_-dicarboxylic acids, which enabled the coexistence of the Cit^−^ and Cit^+^ ecotypes [17].

Beginning at 44,000 generations, however, we were unable to detect Cit^−^ cells in the population. Here we document that the Cit^−^ ecotype went extinct in the population. Prior to that time, the Cit^−^ lineage was present at frequencies of one or more percent of the total population, which had increased to >10^8^ cells per mL (and >10^9^ cells in the 10-mL of medium in the flask) at the end of the daily growth cycle. Even after the 100-fold daily dilution, a lineage that comprised 1% of the total population would include many thousands of cells. Therefore, the extinction of the Cit^−^ ecotype cannot be explained by demographic stochasticity (i.e., as a statistical fluctuation associated with a simple birth-death process). We therefore hypothesized that the extinction was caused by the on-going evolutionary adaptation of the Cit^+^ lineage. Given the larger population size and higher mutation rate of the Cit^+^ lineage, such an outcome would not be unexpected. There are multiple ecophysiological scenarios whereby the Cit^+^ lineage could have evolved to drive the Cit^−^ ecotype extinct. If the Cit^+^ bacteria had evolved to export fewer or less valuable C_4_-dicaboxylate molecules, then this would have reduced the carbon and energy available to the Cit^−^ cells. Similarly, if the Cit^+^ bacteria, or a subpopulation within the Cit^+^ clade, had evolved improved ability to compete for either the glucose or the C_4_-dicaboxylates, then that shift also could have driven the Cit^−^ ecotype extinct.

Beneficial mutations in the LTEE require hundreds, if not thousands, of generations to achieve fixation, with progressively longer periods required in later generations as the opportunity for further fitness improvement declines (Wiser et al. 2013). Therefore, if the hypothesis that the on-going adaptation of the Cit^+^ lineage drove the Cit^−^ lineage extinct was correct, then we would expect the responsible Cit^+^ mutants to have been well-established numerically (even if still a minority genotype) long before the extinction played out; and we would therefore also expect the extinction to be highly reproducible if we replayed evolution from the sample taken within 500 generations before the extinction of the Cit^−^ ecotype.

We tested whether the extinction of the Cit^−^ lineage was deterministic, in the sense of being repeatable going forward from the 43,500-generation sample, by (i) reviving that population sample along with an earlier one from generation 40,000; (ii) propagating 20 replicate populations derived from each of those samples for 500 generations; and (iii) determining whether the Cit^−^ ecotype was present after each of the 40 total replays. Based on the considerations above, we predicted that the Cit^−^ type would persist in most or all of the replay populations started from the 40,000-generation sample, whereas the Cit^−^ lineage would go extinct in most or all of the replays started with the 43,500-generation sample. We also tested whether a Cit^−^ clone could invade and thereby re-establish its frequency-dependent coexistence with Cit^+^ clones and populations from after the extinction event.

## Methods

### *Long-term evolution experiment with* E. coli

The LTEE is an ongoing experiment in which 12 independent populations of *E. coli* are evolving in a glucose-limited Davis-Mingioli minimal medium, called DM25. Six of the populations were founded from REL606, a clone that cannot grow on arabinose (Ara^−^) and six were founded from REL607, a clone that differs from REL606 by a point mutation that confers the ability to grow on arabinose (Ara^+^). DM25 does not contain arabinose, and REL606 and REL607 have equal fitness in the LTEE environment. The focal population in this chapter, Ara-3, was founded from REL606 and is Ara^*−*^. In addition to having 25 ml/L glucose, DM25 contains 326.6 mg/L citrate, an additional carbon source that *E. coli* cannot consume under aerobic conditions. However, a lineage in Ara-3 evolved the ability to consume citrate after more than 30,000 generations [14]. The Cit^+^ lineage coexisted with a Cit^−^ lineage for at least 10,000 generations. Cit^+^ and Cit^−^ clones used in our experiments were identified as described in Blount et al. (2012). More details on the LTEE methods are described in Lenski et al. (1991).

### Assays for presence of the Cit^−^ phenotype and genotype

We tested for the extinction of the Cit^−^ lineage using two methods. First, we measured the frequency of cells with the Cit^−^ phenotype in both DM25, the standard culture medium of the LTEE, and DM25 without citrate. We used DM25 without citrate in addition to the standard culture medium because the proportion of Cit^−^ cells is higher when populations are grown in medium without citrate, thus increasing the probability of detecting them. We knew from prior experiments that Cit^−^ cells were present in the population through at least 43,000 generations [17], and so we revived frozen population samples at 1,000 generation intervals from 40,000 to 46,000 generations as well as populations from 43,500 and 50,000 generations. Revived samples were grown overnight in Luria-Bertani medium (LB), then diluted 10,000-fold into DM25 or DM25 without citrate and grown for 24 h. The cultures were then diluted 100-fold into the same two media and grown again for 24 h. We diluted and plated samples from each culture onto tetrazolium arabinose (TA) indicator agar plates [13, 20]. From the TA plates, we tested 64 colonies from each culture for growth on minimal citrate plates [14]. For any colony that did not grow on minimal citrate, we confirmed the Cit^−^ phenotype using Christensen’s agar [21], which produces a color reaction in response to growth on citrate.

We tested for the presence of the Cit^−^ genotype in populations from the same time points using the polymerase chain reaction (PCR) to amplify a segment of the genome known to have been deleted in the Cit^+^ clade [18]. In addition to serving as a confirmation of the results from the phenotypic assay, the PCR assay also tested the possibilities that (i) the Cit^−^ phenotype was still present in the population, but had lost the ability to form colonies, and (ii) the Cit^−^ clade was still present but had evolved the ability to consume citrate.

### Replays of the extinction event

We hypothesized that ongoing adaptation of the Cit^+^ lineage caused the extinction of the Cit^−^ lineage. If this were the case, then the Cit^+^ genotypes that drove the Cit^−^ lineage extinct should already have been present in the population sample collected and frozen within the 500 generations prior to extinction, given the time required for new beneficial mutations to fix in an LTEE population. To test whether adaptation of the Cit^+^ lineage had, in fact, caused the Cit^−^ extinction, we conducted evolutionary replay experiments with the samples from 40,000 and 43,500 generations of population Ara-3. Based on our hypothesis, we expected Cit^−^ to go extinct in many or all of the replay populations founded from the 43,500-generation sample, but no or few extinctions in replay populations founded from the 40,000-generation sample. We maintained 20 replicate populations started from each sample for 500 generations under the same conditions as the LTEE. Samples from these populations were frozen every 100 generations. We then tested for the presence of the Cit^−^ lineage in the final, 500-generation samples by the PCR assay.

To test for Cit^−^ extinction over a longer time period, we also propagated one population revived from the 43,000-generation sample of population Ara-3 for 2,500 generations. Samples of this population were frozen every 500 generations. We tested for the presence of the Cit^−^ lineage by the PCR assay at each 500-generation interval.

### Invasion experiments

If our hypothesis that ongoing adaptation of the Cit^+^ lineage caused the extinction of the Cit^−^ lineage were correct, then Cit^−^ cells should not be able to reinvade the Cit^+^ lineage after the extinction. We considered two possible cases of adaptation of the Cit^+^ lineage. In one, the trait that allowed the Cit^+^ lineage to exclude the Cit^−^ lineage spread to fixation or near fixation in the Cit^+^ clade. In that case, we would expect that a Cit^−^ invader could not reinvade cultures of either individual Cit^+^ clones or entire Cit^+^ populations from after the extinction event. Alternatively, the Cit^+^ trait that led to exclusion of Cit^−^ might have spread only into a subpopulation of the Cit^+^ clade, perhaps because the fitness benefit of the trait was frequency dependent or perhaps because its benefit was small and therefore slow to spread. If this were the case, then we would expect that a Cit^−^ invader could reinvade cultures of at least some Cit^+^ clones, but not Cit^+^ populations from after the extinction event.

To test whether the extinction of the Cit^−^ lineage was caused by changes in the Cit^+^ lineage that eliminated the Cit^−^ ecotype’s advantage when rare, we tracked whether a Cit^−^ clone could become established when introduced at low and high frequencies to cultures of Cit^+^ bacteria after the extinction event. To monitor the abundance of the Cit^−^ clone in the population, we used a selectively neutral Ara^+^ mutant of a Cit^−^ clone from 43,000 generations as the invading strain. Because the Ara-3 population is Ara^−^, the Cit^+^ bacteria cannot grow on citrate-free minimal arabinose plates (same recipe as minimal citrate, but with 4 g/L L-arabinose in place of citrate), allowing us to track the population size of the initially rare Cit^−^ clone.

The invading Cit^−^ clone was added to three replicate cultures of Cit^+^ clones from 40,000, 42,000, 43,000, 44,000, 45,000 and 50,000 generations at initial densities of ∼3 × 10^3^ (low) and ∼3 × 10^8^ cells/mL (high). We included time points before the extinction as controls, because we expected that the Cit^−^ clone could invade those Cit^+^ clones. In addition, we added the same Cit^−^ strain to whole-population samples from generations 44,000, 45,000, and 50,000. We did not include whole-population samples from time points before the extinction, because the Cit^−^ cells already present in those samples would confound the assay. In all cases, the clones and populations were revived from frozen stocks and acclimated in DM25 for two days prior to the start of the experiment, as described for the experiments above. Each day for 7 days, we transferred the cultures via 1:100 dilutions and maintained them under the same conditions as the LTEE. Each day we measured the Cit^−^ population size by plating on citrate-free minimal arabinose agar. All reported cell densities are based on colony-forming units.

### Data availability

Data will be deposited to the Dryad digital repository upon acceptance.

## Results

### Assays for presence of the Cit^−^ genotype and phenotype

The PCR assay for the presence of the Cit^−^ genotype produced clear bands at all time points up to and including 43,500 generations (Fig 1). At 44,000 generations and beyond, we did not observe any amplification. Examination of phenotypes also indicated that the Cit^−^ lineage went extinct between 43,500 and 44,000 generations. Cit^−^ cells were present through 43,500 generations in all cultures, but they were not detected at any later time point. Prior to the 44,000-generation sample, on average 4 of the tested 64 colonies (∼6%) had the Cit^−^ phenotype when cultures were grown in DM25 medium (Fig 1). Note that this proportion does not necessarily represent the proportion of Cit^−^ bacteria in the population because plating efficiency (the proportion of cells that form colonies) might vary between lineages and over time. We also tested the Cit^+^/Cit^−^ phenotype of colonies produced by cells from cultures that were grown in DM25 without citrate, a medium that selects for Cit^−^ bacteria in the population. In DM25 without citrate, 41 of 64 tested colonies (∼64%) had the Cit^−^ phenotype, on average, prior to the extinction. There was no indication of a downward trend in the proportion of Cit^−^ bacteria in the generations before the extinction.

**Fig 1:**
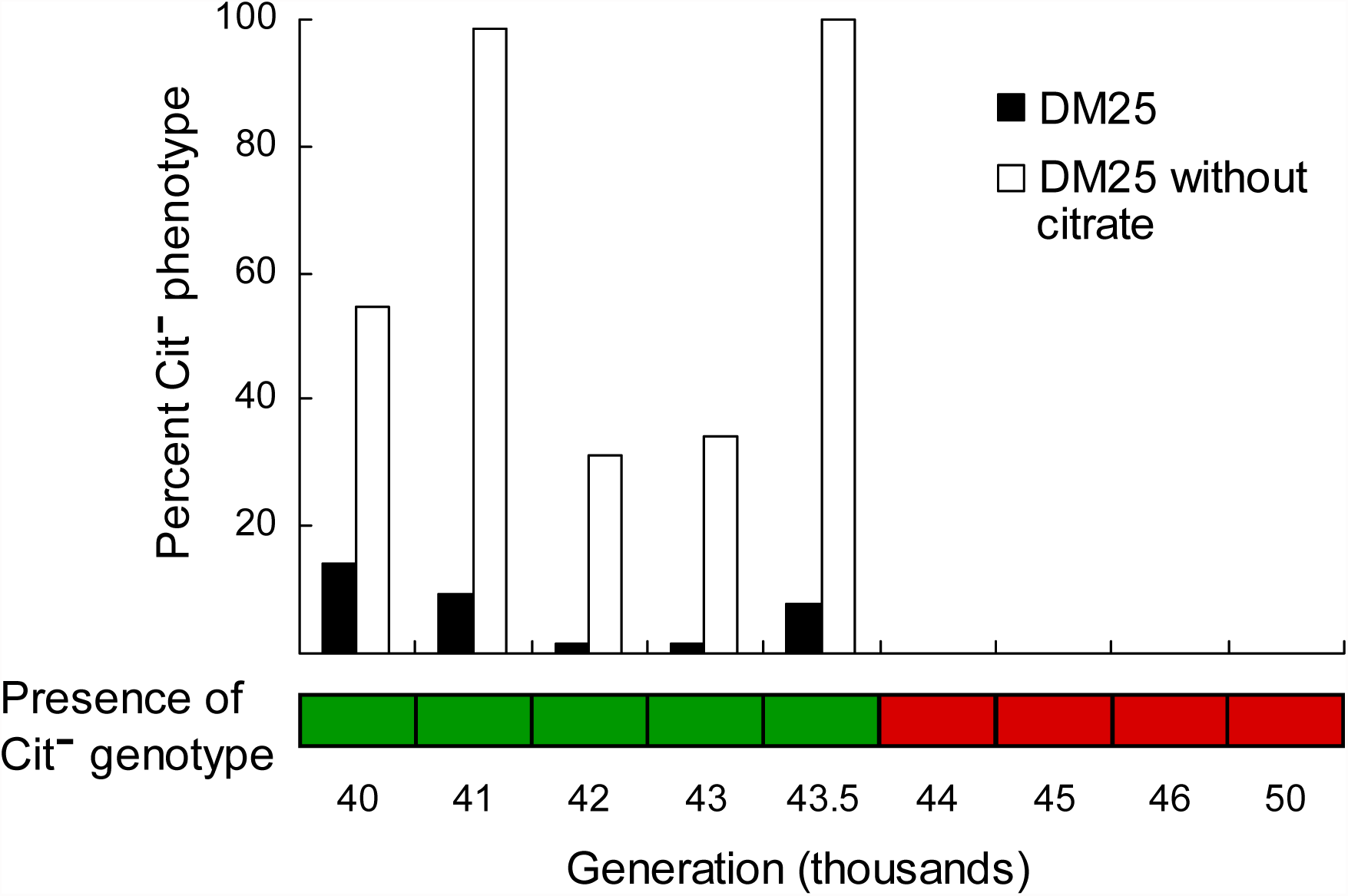
Recovery of Cit^−^ bacteria by generation from the Ara-3 population of the LTEE. The bar graph shows the percentage of clones with a Cit^−^ phenotype recovered in two media, DM25 and DM25 without citrate. The lower grid indicates the presence (green) or absence (red) of the Cit^−^ genotype, as determined by a PCR assay. Taken together, these data show that the Cit^−^ lineage went extinct between 43,500 and 44,000 generations.

### Replays of the extinction event

Twenty replicate populations founded from the 43,500-generation population sample evolved for 500 generations under the standard LTEE conditions. PCR assays showed that the Cit^−^ clade persisted in all 20 cases through the end of the experiment. The Cit^−^ lineage also persisted in all 20 of the control populations founded from the 40,000-generation population sample as well as in the population founded from the 43,000-generation population sample that evolved for 2,500 generations.

### Invasion experiments

When a 43,000-generation Cit^−^ strain was introduced at a low density (∼3 × 10^3^ cells per mL) to a culture containing a single Cit^+^ genotype from several time points before and after the extinction, the population of the Cit^−^ clone increased by about 1000-fold, reaching an average density of ∼4 × 10^6^ cells per mL after 7 days (Fig 2A). When the same Cit^−^ clone was added at a high density (∼3 × 10^7^ cells per mL), the Cit^−^ population declined to the same average population size of ∼4 × 10^6^ cells per mL on day 7. These results indicate that Cit^−^ and Cit^+^ clones can coexist owing to a negative frequency-dependent interaction, regardless of whether the Cit^+^ clone was isolated before or after the extinction of the Cit^−^ lineage.

**Fig 2:**
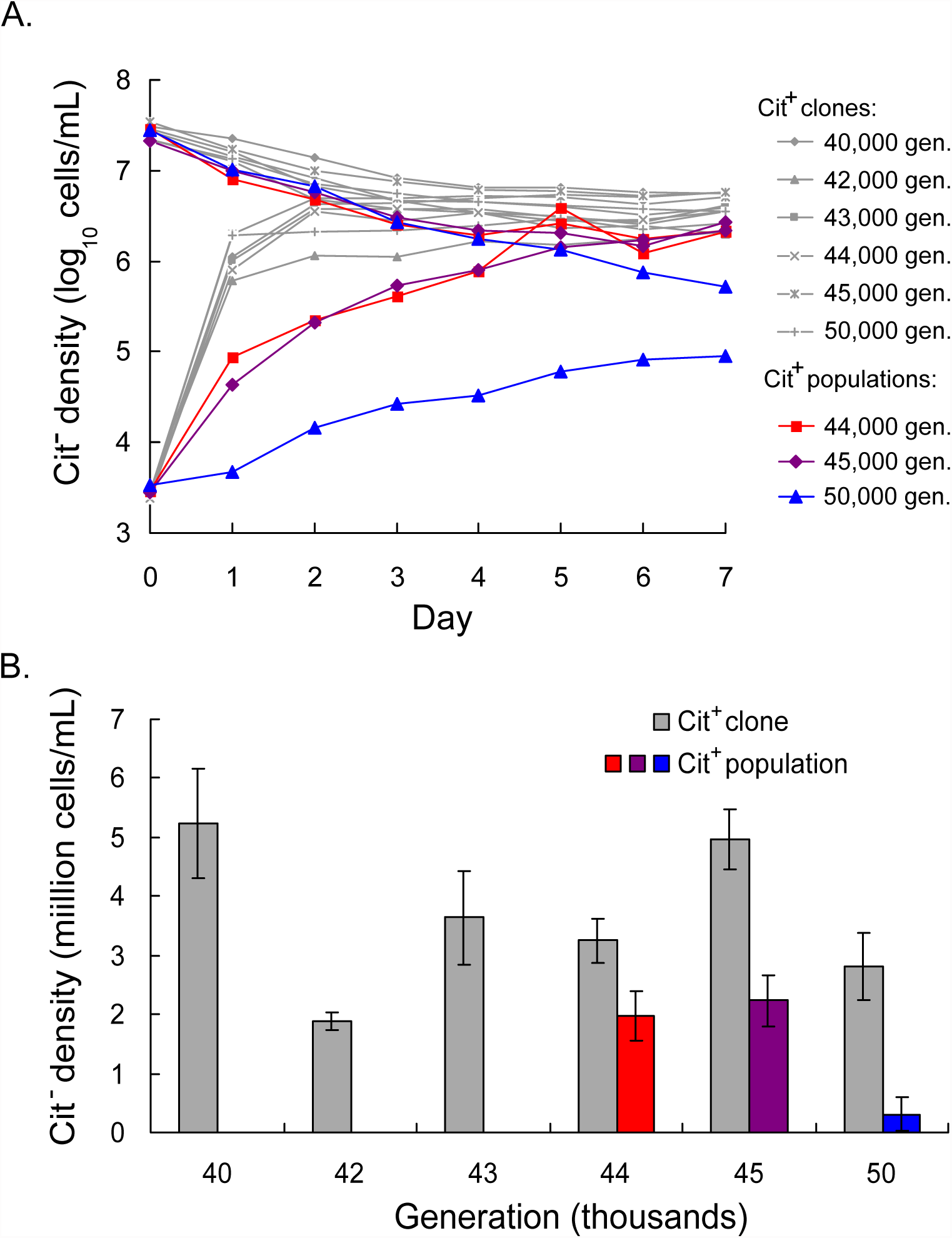
A Cit^−^ clone can reinvade the Ara-3 population after the extinction event. A. Trajectory of the mean population density of the Cit^−^ clone over 7 days, starting from both high and low initial densities and introduced into Cit^+^ clones or populations as indicated in the legend at right. Error bars are omitted for clarity. B. Mean density of the Cit^−^ clone on day 7 of the invasion experiments, including both high and low initial density treatments. The generation of the resident Cit^+^ clone (gray bars) or population (colored bars) is shown along the x-axis. Error bars are 95% confidence intervals.

We also observed invasions and coexistence mediated by negative frequency-dependent selection when the same Cit^−^ clone was added to cultures containing the complete mixture of Cit^+^ genotypes present in the population at 44,000, 45,000, and 50,000 generations. When using the 44,000- and 45,000-generation populations, the final density of the Cit^−^ clone was within the general range observed when it invaded the Cit^+^ clones, although its initial rate of increase when rare was noticeably slower (Fig 2A). The equilibrium density of the Cit^−^ clone was also slightly lower (Fig 2B) when it invaded the entire population than when it invaded a clone from the same generation. When added to the 50,000-generation population—more than 6,000 generations after the extinction of the Cit^−^ lineage—the 43,000-generation Cit^−^ clone could still invade the Cit^+^ population (Fig 2A). However, the final Cit^−^ population size was an order of magnitude lower (3 × 10^5^ cells per mL, mean of high and low initial density treatments) than the final density of Cit^−^ cells in any other treatment (Fig 2B).

## Discussion

Three lines of evidence indicate that the Cit^−^ ecotype, which had coexisted with the Cit^+^ ecotype for more than 10,000 generations, went extinct between 43,500 and 44,000 generations. First, no clones with the Cit^−^ phenotype were isolated from 44,000 generations or later (Fig 1), indicating that the Cit^−^ phenotype was absent from (or at least much rarer in) the population. Second, PCR assays also failed to amplify a Cit^−^ lineage-specific locus from population samples collected after 43,500 generations. Third, if the Cit^−^ ecotype remained in the population, then the Cit^−^ ecological niche would be occupied and the Cit^−^ bacteria would be at a frequency-dependent equilibrium with Cit^+^. In this case, a Cit^−^ strain from an earlier generation, when added at low density, should not be able to invade the population. In contrast to this prediction, however, a Cit^−^ clone from 43,000 generations reinvaded the Ara-3 populations at multiple time points tested after the Cit^−^ extinction, including even 6,000 generations later (Fig 2).

The ability of a Cit^−^ clone to reinvade the population long after the Cit^−^ ecotype went extinct also implied that, contrary to our predictions, the Cit^−^ extinction was not caused by the Cit^+^ bacteria evolving to exclude the Cit^−^ ecotype from the population. Our evolution replay experiments also supported this result. The Cit^−^ lineage persisted for 500 generations in all 20 replicate populations founded from the last population sample prior to the Cit^−^ extinction, as well as in 20 replicate populations started from an earlier point. Moreover, in an additional population that was founded from the 43,000-generation population, the Cit^−^ lineage has continued to coexist with the Cit^+^ ecotype through at least generation 45,500, which is more than 1,500 generations after the Cit^−^ lineage went extinct. The Cit^−^ extinction in Ara-3 therefore appears to be a rare occurrence, and not a repeatable outcome of competitive exclusion by Cit^+^.

As explained in the Introduction, we expected that, if the extinction of the Cit^−^ lineage had been driven by on-going adaptive evolution of the Cit^+^ lineage, then that adaptation would be captured in the population sample collected immediately prior to extinction. However, our results also eliminate the possibility that the extinction of the Cit^−^ lineage was caused by some rare, highly beneficial mutation that arose in the Cit^+^ lineage at some point after the prior population sample and spread quickly through the population. The presence of such a mutation in the Cit^+^ population would have prevented the reinvasion of the population by Cit^−^ cells in subsequent generations.

Thus, the cause of the Cit^−^ extinction remains unknown. We have rejected our hypothesis that evolutionary changes in the Cit^+^ bacteria drove the extinction. Moreover, a purely stochastic extinction of the Cit^−^ lineage is also extremely unlikely. The LTEE is propagated by 1:100 daily transfers, and the total number of Cit^−^ cells in the culture prior to transfer (i.e., after 24 h) was consistently measured at >10^7^ prior to their extinction. Therefore, the probability that no Cit^−^ cells would be present in the population after a transfer, given that each cell has a 99% chance of not being transferred, is less than 0.99^10^,^000^,^000^ ≈ 10^-43^,^700^. Even if some perturbation were to cause a moderate decline in the Cit^−^ population size, its density should then have tended to increase back towards its equilibrium owing to the negative frequency-dependent relationship between the Cit^+^ and Cit^−^ ecotypes [14, 17], making a stochastic extinction even more unlikely.

In populations large enough that demographic stochasticity is unlikely to cause extinction, the most likely causes of random extinction are environmental stochasticity and random catastrophes [3]. Environmental stochasticity refers to changes in population size caused by environmental variation over time, which should be minimal in the context of a controlled laboratory experiment like the LTEE. Furthermore, the large size of the Cit^−^ population should have made it robust to environmental stochasticity caused by the typical fluctuations that simply cannot be avoided in any experimental setting. We propose, therefore, that the most likely cause of the Cit^−^ extinction was a “random catastrophe”, that is a large, rare event that reduced the size of the Ara-3 population as a whole or, at least, the Cit^−^ lineage within it. We do not know the nature of this random catastrophe, but it could have been some unusual variation in laboratory conditions, such as soap residue in a flask or problems with the deionized water used to make the culture medium. Although the Cit^−^ population was too large for purely stochastic extinction from demographic fluctuations, its population size was substantially smaller than that of the Cit^+^ population, making the Cit^−^ lineage more vulnerable to such occurrences [22, 23].

In addition to demonstrating that the extinction of the Cit^−^ population was not a deterministic event, we also found evidence that there has been ecological diversification in the Cit^+^ clade. The equilibrium Cit^−^ population density was significantly lower when it was grown with the whole Cit^+^ population than when it was grown with a single Cit^+^ clone at each time point tested. This result suggests that some Cit^+^ genotypes in the population were better at competing with the Cit^−^ invader for shared resources than were the particular Cit^+^ clones tested individually at those time points. The effect of this diversity was particularly pronounced at 50,000 generations, where the equilibrium population size of the Cit^−^ cells when grown together with the Cit^+^ population was nearly an order of magnitude lower than in earlier generations (Fig 2). This decline in the reinvasion ability of the Cit^−^ clone indicates that, by 50,000 generations, a subpopulation of Cit^+^ bacteria had evolved improvements that exploited or otherwise impacted the niche previously occupied by the Cit^−^ bacteria. Thus, rather than the Cit^−^ lineage having been driven extinct by the Cit^+^ bacteria evolving into its niche, it instead appears that the extinction of the Cit^−^ lineage opened a niche that a Cit^+^ subpopulation has begun to exploit.

The occurrence of this extinction during an evolution experiment allowed us to test the hypothesis that the extinction was caused by the evolution of other organisms, something that would not be possible to test in natural systems. Having found that the extinction of the Cit^−^ lineage was likely caused by a rare environmental perturbation—a random catastrophe, as it were—we can take advantage of another feature of experimental evolution. In particular, we can now study, going forward, the consequences of that extinction event by following two ‘alternative histories’ of evolution, one with and one without the extinction. To that end, we are continuing to propagate the population that was founded from a sample taken before the Cit^−^ extinction and evolved for 2,500 generations. We have designated it Ara-7, and it is the thirteenth population in the LTEE, although it is well behind the other populations in terms of its total elapsed generations owing to the fact that it was only re-started from the 43,000-generation sample of Ara-3 when that population was at generation 56,500. In the future, Ara-3 and Ara-7 together will allow us to compare the outcomes in these two populations, which differ in the occurrence of the extinction of the Cit^−^ bacteria. Will the continued presence of the Cit^−^ cells in Ara-7 prevent members of the Cit^+^ clade from diversifying and occupying the Cit^−^ niche, as the Cit^+^ clade appears to be doing in Ara-3? Will the Cit^+^ bacteria in Ara-3 eventually evolve to occupy fully the Cit^−^ niche, such that pre-extinction members of the Cit^−^ clade can no longer re-invade the Cit^+^ population? Might a new clade with a Cit^−^ phenotype emerge from within the Cit^+^ clade in Ara-3? All this remains to be seen.

## Acknowledgments

We thank Neerja Hajela, Daniel Mitchell, Maia Rowles, and Jennifer Jimenez for laboratory assistance. This research was supported, in part, by an EPA STAR Fellowship and a MSU Distinguished Graduate Student Fellowship to C.T., an NSF grant (DEB-1019989) to R.E.L., a grant from the John Templeton Foundation to R.E.L. and Z.D.B., and the BEACON Center for the Study of Evolution in Action (NSF Cooperative Agreement DBI-0939454).

